# Mitotic lineage adds predictive information beyond cell type in the *C. elegans* connectome

**DOI:** 10.1101/2022.11.01.514680

**Authors:** Jordan K. Matelsky, Brock Wester, Konrad P. Körding

## Abstract

During nervous system development, repeated cell divisions of the zygote give rise to a “family tree” of neurons related by mitotic lineage. The developmental process also gives rise to neuronal connections, and neurons phenotypically converge to different cell types. Neural connections are steered by cell type, but they may be driven in part by lineage: We do not know if lineage matters for the developmental neurogenesis process. We thus asked if mitotic lineage predicts neural connections beyond cell type alone. Using three *C. elegans* datasets, we fit models for edge prediction tasks: predicting synaptic targets, and predicting synaptic sources. Adding the mitotic lineage improved these predictions. Our results suggest that the family tree matters for the connections made by neurons, and that developmental lineage is a variable that should be considered more deeply in models of connectome assembly.

## Introduction

Brains are built developmentally: cells divide to make new cells, and repeated mitotic branching yields a lineage tree. That tree captures shared developmental ancestry. We ask if access to lineage information improves models of neuronal connectivity beyond what is possible with cell type alone (**Fig. 1**). If lineage matters, then neuroscience should consider this developmental variable much more deeply when modeling how nervous systems are assembled.

**Figure 1.**
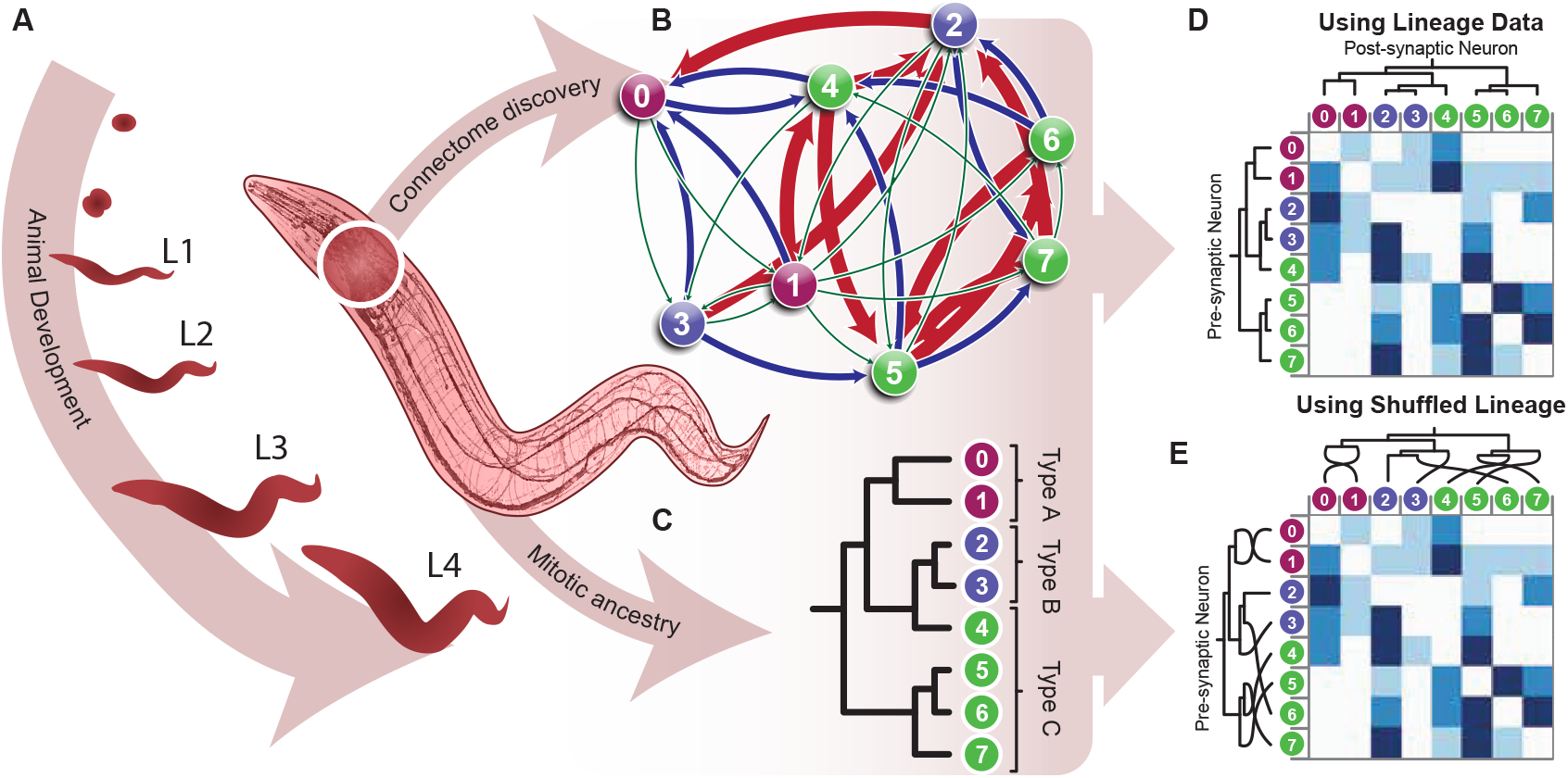
Conceptual framing for lineage-connectome integration. **A**, development yields a mitotic lineage tree. **B**, connectomics yields a synapse-resolved directed graph. **C**, cell-type and lineage labels can be aligned to graph nodes. **D**, pairwise and motif-level analyses relate lineage structure to connectivity structure. **E**, lineage-shuffle null logic asks whether the lineage-to-connectome mapping carries signal beyond the fixed tree and graph topologies. Cell and animal morphology graphics are based on OpenWorm and BossDB resources (***Szigeti et al., 2014; Hider et al., 2019***)

Connectome assembly is generally framed as a developmental process in which neurogenesis, neurite outgrowth, and local partner-recognition rules progressively constrain which cells can meet and synapse (***Kerstjens et al., 2022***). In *C. elegans*, this question is especially tractable because neurite neighborhoods remain broadly stable across development even as synaptic connectivity matures (***Moyle et al., 2021; Witvliet et al., 2021***). Cell type is a central organizing concept in neuroscience, but it does not fully specify how connectomes are assembled (***Thompson et al., 2014; Erö et al., 2018; Zeng and Sanes, 2017; Barabási and Czégel, 2021***). Physical geometry also matters, yet geometric proximity is likewise an incomplete predictor of connectivity (***Varshney et al., 2011***). The *C. elegans* system enables a direct test because complete lineage resources and synapse-resolved connectomes are available across development and in a mature reference (***Bhatla, 2011; Witvliet et al., 2021; Cook et al., 2019***). **Figure 2** shows this alignment directly for the `Witvliet_8` dataset: the lineage tree, cell-type labels, and connectome adjacency can be coregistered.

**Figure 2.**
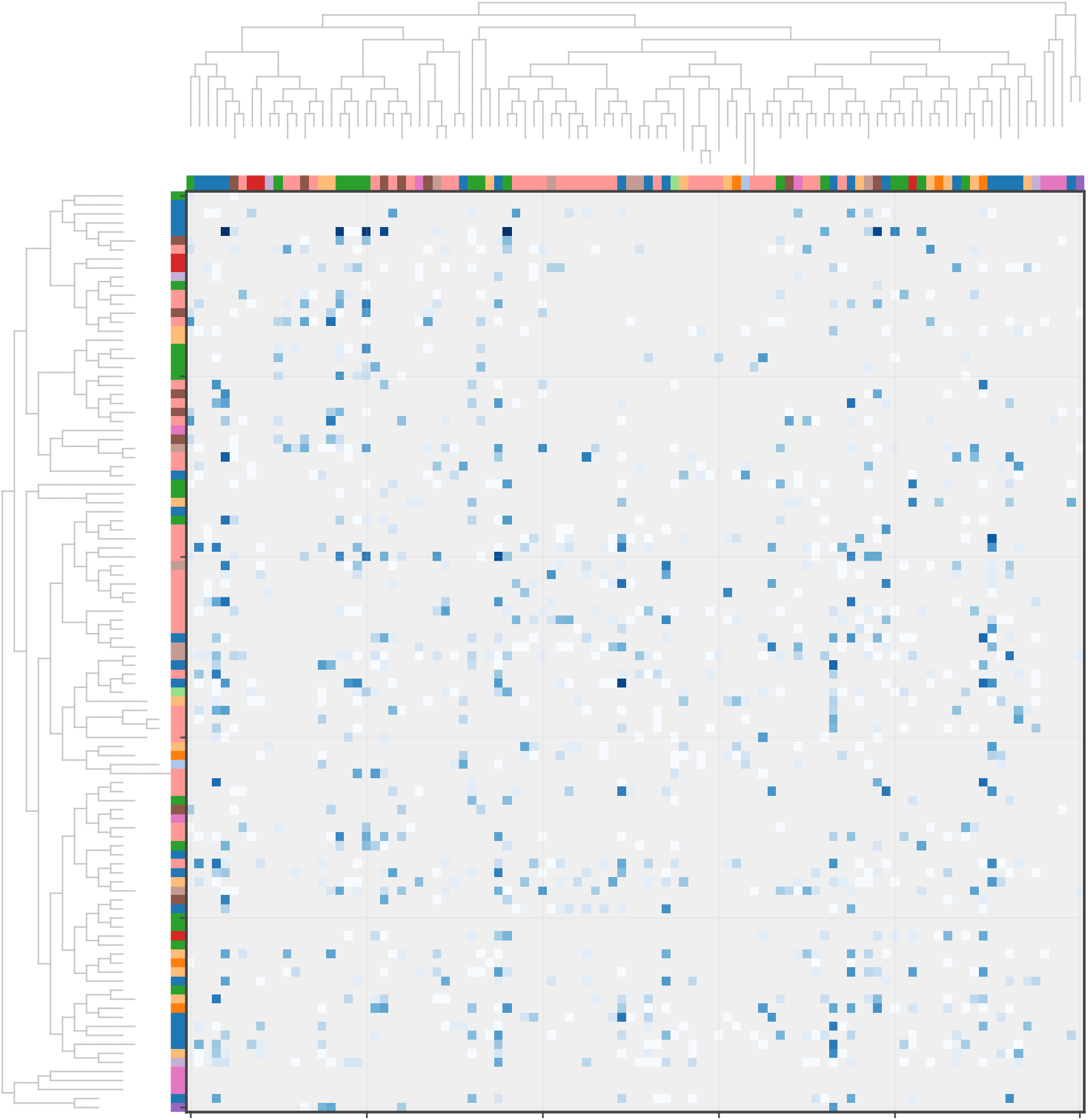
C. elegans lineage aligned to the connectome. Leaves of the two (identical) dendrograms along the perimeter of the plot correspond to neurons shared between the `Witvliet_8` connectome and the lineage tree, colored by composite *cell type* label. The same lineage-derived leaf order is used for the rows and columns of the directed chemical-synapse adjacency matrix, with local branch order chosen to keep nearby connectivity profiles together.

Here we focus on the late developmental and mature regime, analyzing `Witvliet_7, Witvliet_8`, (***Witvliet et al., 2021***) and the Cook/White N2U mosaic (N2U, ***Cook et al***. (***2019***)) with held-out linear models for two directed edge-prediction tasks: predicting a source neuron’s targets and predicting a target neuron’s sources (**Fig. 4**). Each ordered row is a candidate directed connection *u* → *v*. In target prediction, we hold the source neuron *u* fixed and ask which candidate targets *v* it connects to; in source prediction, we hold the target neuron *v* fixed and ask which candidate sources *u* connect into it. Here, held-out prediction is used as an assay of information content rather than as an attempt to maximize predictive accuracy; the question is whether lineage adds out-of-sample signal beyond cell type.

## Results

### Cell type is already a strong predictor of directed connectivity

Before asking whether lineage helps, we first establish what a fair non-lineage baseline already captures. After all, cell type is known to be a strong predictor of connectivity. Using cell-type descriptors alone, the directed models already achieved mean held-out test *R*^2^ values of 0.0199 for target prediction and 0.0177 for source prediction (**Fig. 3**). By contrast, a distance-only model sat essentially on the held-out ***R***^2^ = 0 line (Appendix). The interpretation of everything that follows therefore hinges on an incremental question: not whether lineage can predict connectivity from scratch, but whether it improves prediction after cell type has already explained a substantial part of the linear signal.

**Figure 3.**
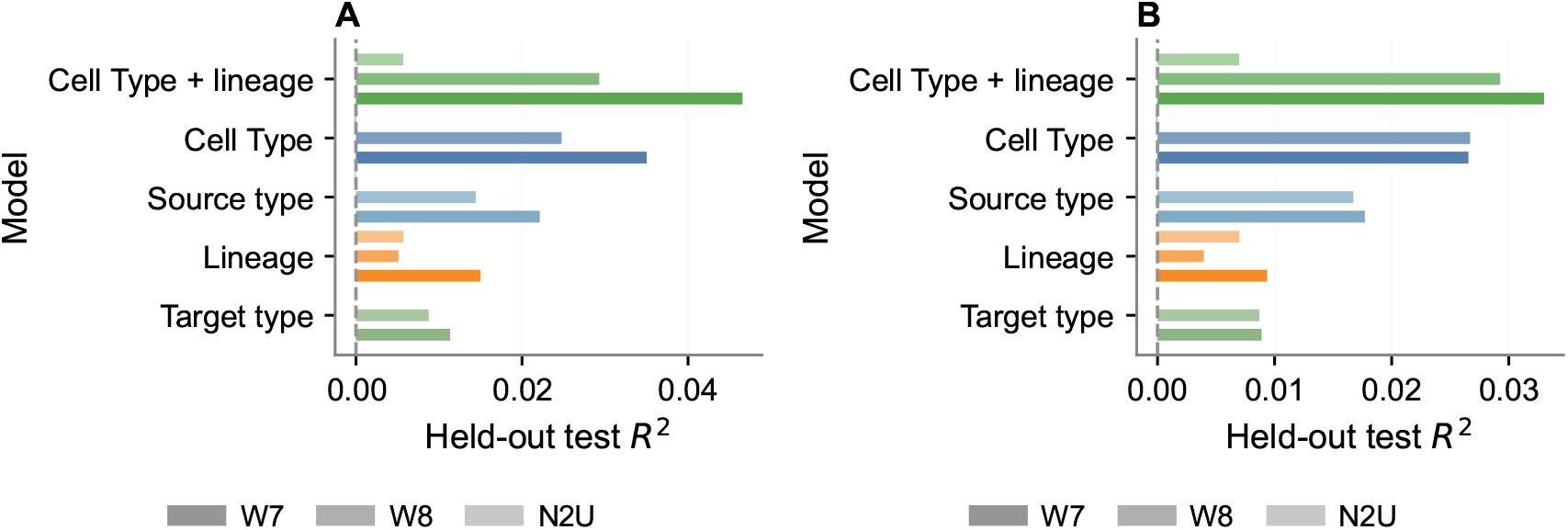
Constituent cell-type baselines. **A**, target-prediction model summaries using cell type descriptors and lineage. B, source-prediction model summaries. Bars show the mean held-out test *R*^2^ across the adult datasets, and error bars show the standard deviation across datasets. This figure establishes the baseline that lineage must beat: source and target type descriptors already explain much of the accessible linear signal, yet adding lineage still improves performance.

### Lineage adds held-out signal beyond cell type in both directed tasks

Adding lineage improved held-out performance in both directed edge-prediction tasks (Table 1; **Fig. 4**). For target prediction (**Fig. 4A**), the mean test *R*^2^ rose from 0.0199 to 0.0272, a gain of 0.0073 (+37%). For source prediction (**Fig. 4B**), it rose from 0.0177 to 0.0231, a gain of 0.0054 (+30%). Crucially, these gains were positive in all three adult datasets.

**Table 1.** Held-out directed-edge predictive gains from adding lineage. The main effect is the increase in held-out test *R*^2^ after adding lineage to the cell-type baseline. Raw model *R*^2^ values are shown for context.

| Task | Cell Type | Cell Type + lineage | $\Delta R^2$ | Relative lift |
| --- | --- | --- | --- | --- |
| Target prediction | 0.0199 | 0.0272 | 0.0073 | +37% |
| Source prediction | 0.0177 | 0.0231 | 0.0054 | +30% |

**Figure 4.**
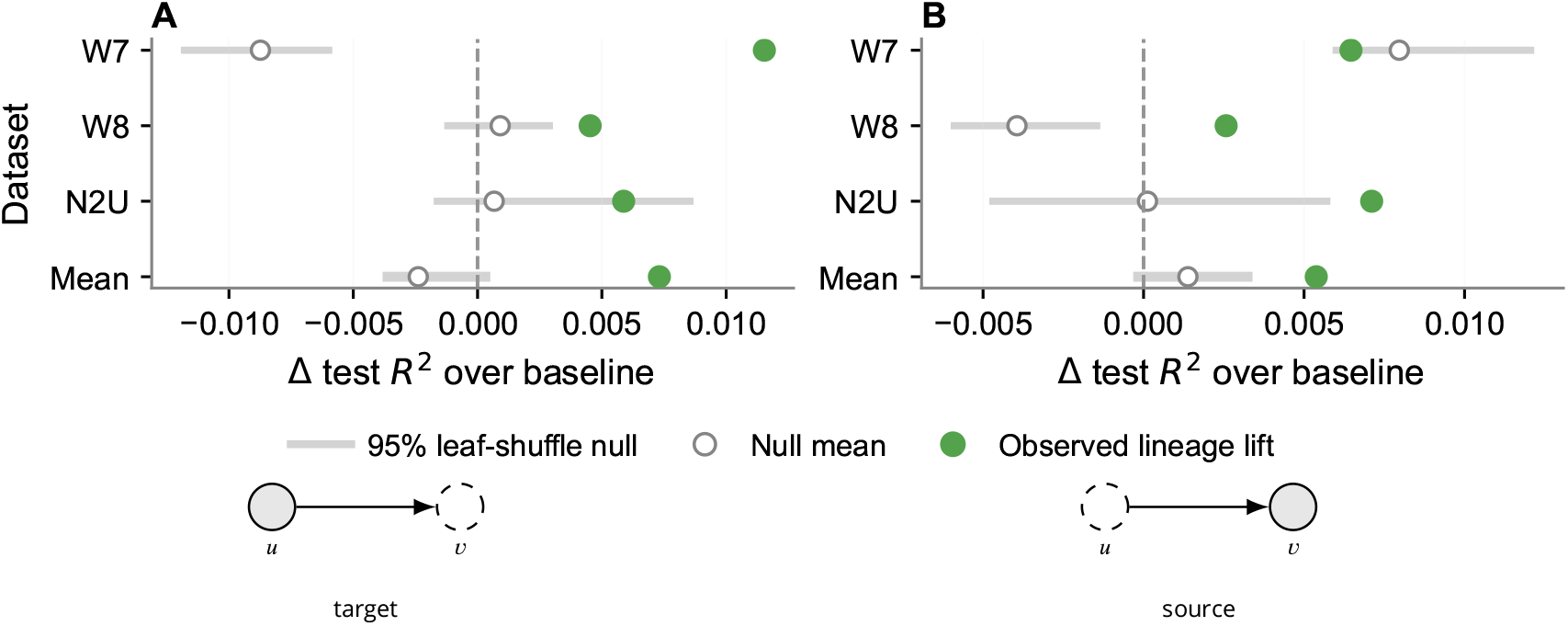
Lineage-leaf-shuffle null. **A**. Target prediction. Target prediction asks whether a fixed source neuron *u* targets a candidate neuron *v*. In schematics, shaded nodes are held fixed within a split group, and dashed-outline nodes denote the candidate partner being predicted. **B**. Source prediction. Source prediction asks the complementary question of whether a candidate neuron *u* is an input to a fixed target neuron *v*. For each dataset, the filled green dot marks the observed held-out gain from adding lineage to the fair baseline, the open circle marks the mean gain after shuffling the mapping between connectome neurons and lineage leaves, and the gray interval shows the central 95% of the shuffled null distribution. The bottom row summarizes the across-dataset mean. Both directed edge-prediction tasks clear the shuffle null.

The lineage-only model performed substantially worse than the combined model in both tasks (0.00860 for target prediction and 0.00677 for source prediction). That pattern is important for interpretation. The result is *not* that lineage is a strong standalone substitute for cell type. (It is not!) Rather, lineage contributes nonredundant information that the cell-type baseline does not already absorb.

### The gain is extinguished with a shuffled-null lineage-to-neuron mapping

If lineage is truly contributing task-relevant information, then the gain should depend on the actual mapping between connectome neurons and lineage leaves, not merely on the fixed shapes of the lineage tree and the connectome. We tested that prediction by permuting the lineage-to-neuron mapping within each dataset (**Fig. 1E**), recomputing lineage features, and refitting the same held-out models (**Fig. 4**). For target prediction, the observed mean gain across datasets was 0.00731 test-*R*^2^ units, whereas the 95% leaf-shuffle null interval for the across-dataset mean ranged from −0.00382 to 0.00051. Source prediction also cleared its null, with an observed mean gain of 0.00538 against a 95% null interval from −0.00032 to 0.00339. This null control sharpens the central claim: the advantage of lineage is not a generic consequence of having some hierarchical feature block, but depends on the real alignment between developmental ancestry and connectome identity.

### Lineage features work best as an ensemble

Having established that lineage adds signal, we next asked which part of the lineage description carries that gain. To do so, we replaced the full lineage block with one lineage term at a time while keeping the same cell-type baseline. Across both directed tasks, the full three-term lineage block performed best overall (**Fig. 5**).

**Figure 5.**
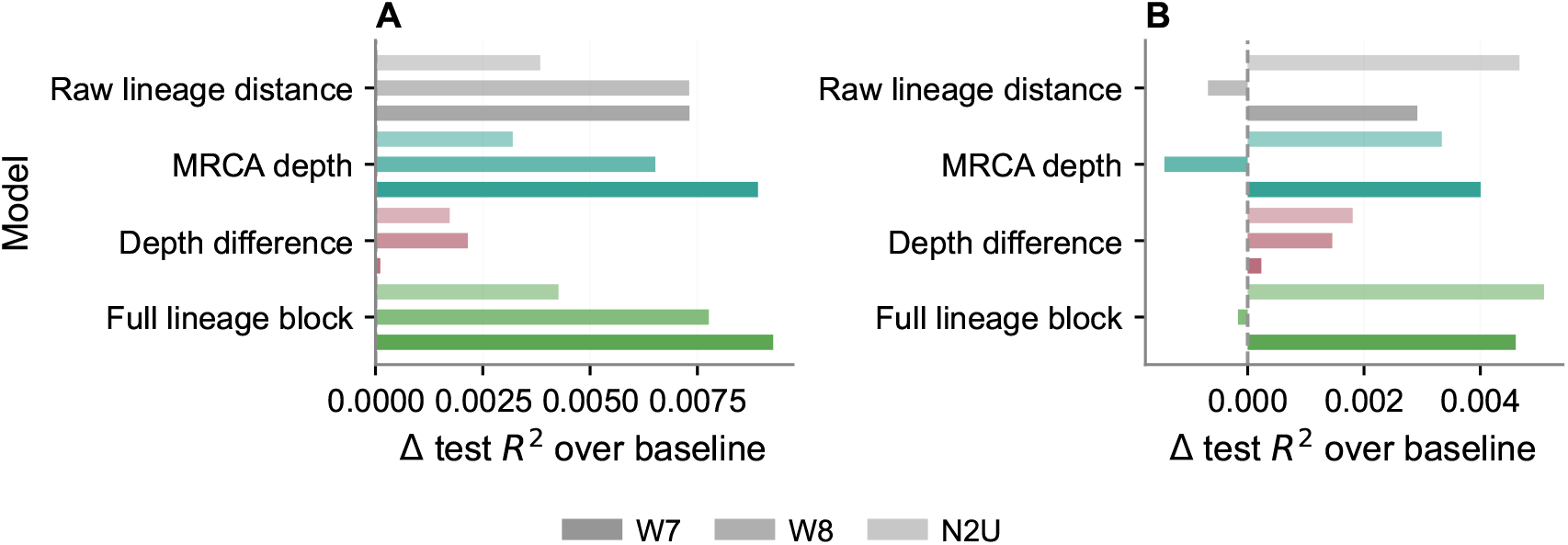
Lineage features work best as an ensemble. **A**, target prediction. **B**, source prediction. For each directed edge-prediction task, bars show the mean held-out improvement in test *R*^2^ over the same cell-type baseline across the three adult datasets, and error bars show the standard deviation across datasets, when adding either raw lineage distance alone, MRCA depth alone, lineage depth difference alone, or the full three-term lineage block. The full block performs best overall, indicating that the lineage effect is distributed across multiple correlated summaries rather than being captured cleanly by any single term.

In target prediction, the full block improved held-out performance by 0.00710 test-*R*^2^ units over cell type alone, slightly exceeding most recent common ancestor (MRCA) depth alone (0.00621) and raw lineage distance alone (0.00615), while lineage-depth difference alone contributed much less. In source prediction, the same full block improved held-out performance by 0.00318, with raw lineage distance alone (0.00230) slightly ahead of MRCA depth alone (0.00197). The interpretation is that no single scalar lineage summary fully captures the effect. Most of the signal is carried by shared developmental ancestry and lineage separation, while absolute depth difference contributes comparatively little.

## Discussion

In late developmental and adult *C. elegans*, mitotic lineage information adds reproducible improvement in predictive models of connectivity beyond cell type. That effect is consistent across datasets and can be eliminated with a lineage-leaf-shuffle null, which argues that the real mapping between lineage and connectome identity carries signal. This is exactly the pattern we expect if developmental ancestry captures a real part of the residual structure left over after cell type has already done much of the predictive work.

The lineage decomposition refines that interpretation. MRCA depth and raw lineage distance carry most of the signal, whereas the absolute difference in lineage depth contributes much less. This suggests that the relevant information is tied more to shared developmental history and lineage separation than to one neuron simply being developmentally “earlier” or “later” than another.

We should not infer from this work that lineage causally determines the connectome. Stronger claims would require richer model classes, additional species, or experimental perturbations that directly test developmental causality. Instead, our results are consistent with the idea that lineage is one of the variables that reduces the residual wiring entropy that must be addressed by neurogenesis.

Genomic-bottleneck arguments suggest that nervous systems cannot specify every synapse independently and instead must compress wiring instructions into reusable developmental programs (***Koulakov et al., 2021***). In that framing, cell type removes part of the uncertainty in connectivity, and our results suggest that developmental ancestry removes a bit more.

Because cell type and lineage are correlated, confound control remains essential (***Pearl, 2009; Angrist and Pischke, 2014***). Here, lineage still provided an improvement after conditioning on type. That illustrates that lineage captures information not already absorbed by cell type, and it argues against the lineage effect being merely a proxy for cell-type misclassification noise.

More broadly, lineage has been linked to neuronal and physiological specialization beyond connectomics (***Li et al., 2012; Cadwell et al., 2020; He et al., 2020***), and emerging lineage-tracing technologies should make cross-system tests increasingly feasible (***Masuyama et al., 2019***). We therefore view lineage-aware models as part of a larger developmental program for understanding how wiring constraints are inherited over cell divisions (***Kerstjens et al., 2022***).

As additional matched reconstructions across individuals and stages become available, it should become possible to estimate lineage effects more precisely and separate reproducible structure from animal-to-animal variation (***Cook et al., 2019; Witvliet et al., 2021***). We therefore view this analysis as a community resource that should grow more informative as the number of public *C. elegans* connectomes increases.

## Methods

### Data, normalization, and folding

Connectomes were accessed through MotifStudio/BossDB endpoints using host IDs `Witvliet_7, Witvliet_8, and N2U` (***Hider et al., 2019; Matelsky et al., 2021, 2025***). These analyses were restricted to late developmental and mature connectomes. Species/dataset metadata were controlled by a YAML registry for reproducibility. Lineage was loaded from WormWeb-derived JSON (***Bhatla, 2011***), and soma locations from OpenWorm-derived coordinates with WormBase-supported identity annotations, with postprocessing performed in Blender (***Szigeti et al., 2014; Harris et al., 2010; Blender Online Community, 2026***). Graph and table analyses were implemented in Python with NetworkX (***Hagberg et al., 2008***).

Neuron identities were aligned by name with normalization rules from the registry. Bilateral neurons were folded with right-side preference; edge weights were aggregated by summation (with mean as an available alternative but unused mode). This folding operation is adopted from prior work in *C. elegans* connectomics (***Witvliet et al., 2021***).

### Prediction tasks

The first task was ordered edge-target prediction, h[*u* → *v*]. Each row corresponds to a candidate source-target pair among neurons shared between the lineage and connectome graphs.

The second task used the same directed edge rows but asked the complementary held-out question of source prediction: can a target neuron’s incoming partners be predicted from candidate sources? The outcome variable remained h[*u* → *v*] but train-test splits were formed within target-neuron groups rather than source-neuron groups.

### Feature blocks

For target prediction and source prediction, the paper’s main baseline block contained decomposed typ indicators for the source and target. A single physical-distance term was evaluated separately as an appendix negative control rather than included in the paper’s main baseline, as it did not substantially improve performance and unnecessarily complicated the models.

The lineage block contained three terms: undirected lineage distance, depth of the most recent common ancestor (MRCA), and the absolute difference between the depths of the two lineage nodes. Additional diagnostic models replaced the full lineage block with one lineage term at a time to determine which component carried the predictive signal.

### Model fitting and evaluation

All models were ordinary least squares fits evaluated on held-out data. For target prediction, train-test splits were formed within source-neuron groups so that each source retained both train and test targets. For source prediction, splits were formed within target-neuron groups. The default held-out fraction was 0.25.

As a negative control, we repeated the combined-model fit after 20 random shuffles of the mapping from overlapping connectome neurons to lineage leaves within each dataset. The lineage tree, connectome, baseline features, and train-test split were held fixed; only the lineage-to-neuron assignment used to compute lineage features changed.

We report held-out test *R*^2^ for each model, but in the Results we emphasize the held-out Δ*R*^2^ gained by adding lineage to the cell-type baseline, because that quantity most directly answers the biological question of interest. We then summarize constituent cell-type models and lineage-component models separately.

### Reproducibility & Rigor

All analyses produce deterministic figures, summary tables, and run manifests. Supplemental materials, including reproducibility details and generated assets, are provided at https://github.com/KordingLab/LineageConnectomics.

## Appendix

### Simple physical-distance terms sit near the held-out *R*^2^ = 0 line

As a simple negative control, we evaluated models that used only the Euclidean distance between source and target somata, as well as models that added this distance term to the cell-type baseline. Across the adult-only directed tasks, distance-only models achieved mean held-out test *R*^2^ values of −0.000084 for target prediction and 0.000082 for source prediction, i.e. essentially on the *R*^2^ = 0 line for this metric. Adding distance to cell type increased mean held-out test *R*^2^ only slightly, from 0.0199 to 0.0217 for target prediction and from 0.0177 to 0.0195 for source prediction (**Fig. 6**). These checks do not imply that geometry is biologically irrelevant; only that this very simple linear distance term is not a strong standalone predictor here.

**Figure 6.**
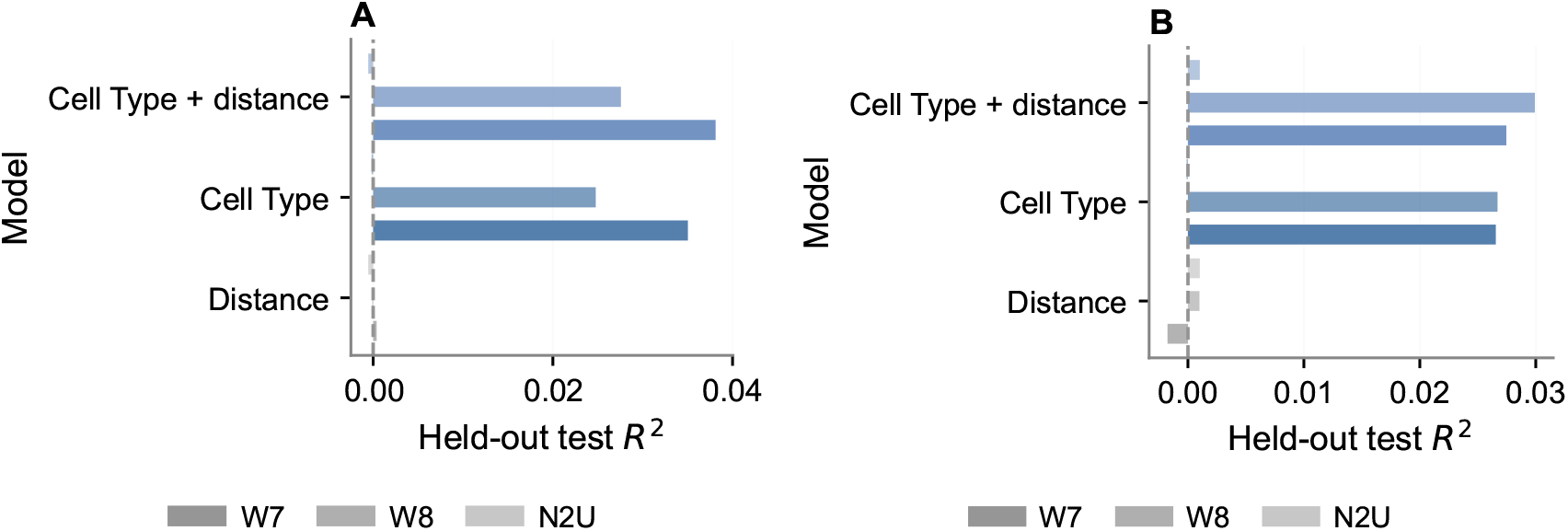
Simple physical-distance terms are weak in this linear setup. **A**, target prediction. **B**, source prediction. Bars show the mean held-out test *R*^2^ across the three adult datasets for distance alone, cell type alone, and cell type plus a single physical-distance term; error bars show the standard deviation across datasets. The dashed vertical line marks *R*^2^ = 0, the constant-predictor reference for this metric. Distance-only models sit at or extremely near that line, while adding distance to cell type changes performance only slightly.

## Supplemental Materials

Supplemental figures, tables, and reproducibility materials are provided at https://github.com/KordingLab/LineageConnectomics. The README documents how to reproduce manuscript outputs from the Snakemake workflow or individual analysis entrypoints.

## Acknowledgements

The authors thank all contributors to *C. elegans* connectomics, and Nikhil Bhatla for curation of the

*C. elegans* lineage tree resources.

